# Intragenomic variability and extended sequence patterns in the mutational signature of ultraviolet light

**DOI:** 10.1101/640722

**Authors:** Markus Lindberg, Martin Boström, Kerryn Elliott, Erik Larsson

## Abstract

Mutational signatures can reveal properties of underlying mutational processes and are important when assessing signals of selection in cancer. Here we describe the sequence characteristics of mutations induced by ultraviolet (UV) light, a major mutagen in several human cancers, in terms of extended (longer than trinucleotide) patterns as well as variability of the signature across chromatin states. Promoter regions display a distinct UV signature with reduced TCG>TTG transitions, and genome-wide mapping of UVB-induced DNA photoproducts (pyrimidine dimers) showed that this may be explained by decreased damage formation at hypomethylated promoter CpG sites. Further, an extended signature model encompassing additional information from longer patterns improves modeling of UV mutation rate, which may enhance discrimination between drivers and passenger events. Our study presents a refined picture of the UV signature and underscores that the characteristics of a single mutational process may vary across the genome.

## INTRODUCTION

Cancer is caused by somatic mutations that alter cell behavior, and the mutational processes that shape tumor genomes are therefore at the core of the disease^1^. Elucidation of such processes and their sequence preferences can provide clinically relevant insights^2,3^ and also allows improved estimation of expected mutation frequencies at recurrently altered positions, which is key when identifying driver events that are under positive selection^4^. A useful approach to discovering and characterizing mutational processes are trinucleotide-based mutational signatures, which describe the probability of mutagenesis at all possible trinucleotide sequence contexts for a given process, normally determined at the whole genome or exome level^5^. However, as cancer sequencing cohorts grow larger, even small errors in our understanding of mutation rate heterogeneity across the genome can lead to false signals of positive selection^6^, motivating an increasingly more detailed understanding of mutational processes and their sequence characteristics.

Specifically in the case of ultraviolet (UV) light, the main mutational process in melanoma and other skin cancers^5^, prior work suggests that the widely adopted trinucleotide model is inadequate. The main basis for UV mutagenesis is the formation of cyclobutane pyrimidine dimers (CPDs) or, at lower frequency, (6,4) photoproducts (6,4-PPs); bulky DNA lesions that bridge neighboring pyrimidines and that may result in C>T or CC>TT transitions^7,8^. The canonical UV mutational signature (“Signature 7”) is consequently dominated by C>T substitutions at dipyrimidine-containing trinucleotides^5^. However, studies of UV-induced DNA damage in melanoma exome data^9^ as well as smaller target templates^10,11^ support that presence of thymines in positions beyond the immediate neighboring bases may confer elevated mutation rates. Additionally, it is known that ETS transcription factor binding site sequences (TTCCG) are associated with strongly elevated CPD formation and mutation rates in melanoma, but only in promoter regions and notably at positions that are variable relative to the motif^12-15^. These intricacies of the underlying mutational process may contribute to recurrent mutations in skin cancers, but cannot be captured using trinucleotide-based UV mutational signatures.

Here, we characterize the sequence signature of somatic mutations arising from UV photoproduct formation, in terms of trinucleotide and extended (beyond trinucleotide) patterns that carry information about mutation rates. We also study variability in these patterns across chromatin states. To gain mechanistic insight, we further generate the first human genome-wide map of CPDs arising from UVB, the main inducer of DNA photolesions in sunlight, which differs in its effects on DNA compared to UVC used in earlier studies. Our results constitute a refined picture of the UV mutational signature and its variability across the genome, with possible implications for the interpretation of recurrent mutations in cancer.

## RESULTS

### A large compendium of predominantly UV-induced somatic SNVs

While most cancer genomes are mosaics of somatic mutations induced by different processes, the study of UV-induced mutations is facilitated by their dominance and abundance in skin cancers. Here, we selected a subset of 130 “high-UV” melanoma whole genomes from an initial set of 221 samples assembled from published studies^16,17^, excluding those with lower burden (<10,000 mutations) or lower fraction UV photoproduct-type mutations (<80 % C>T or CC>TT in a dipyrimidine context; **Fig. 1a** and **Supplementary Table 1**).

**Figure 1.**
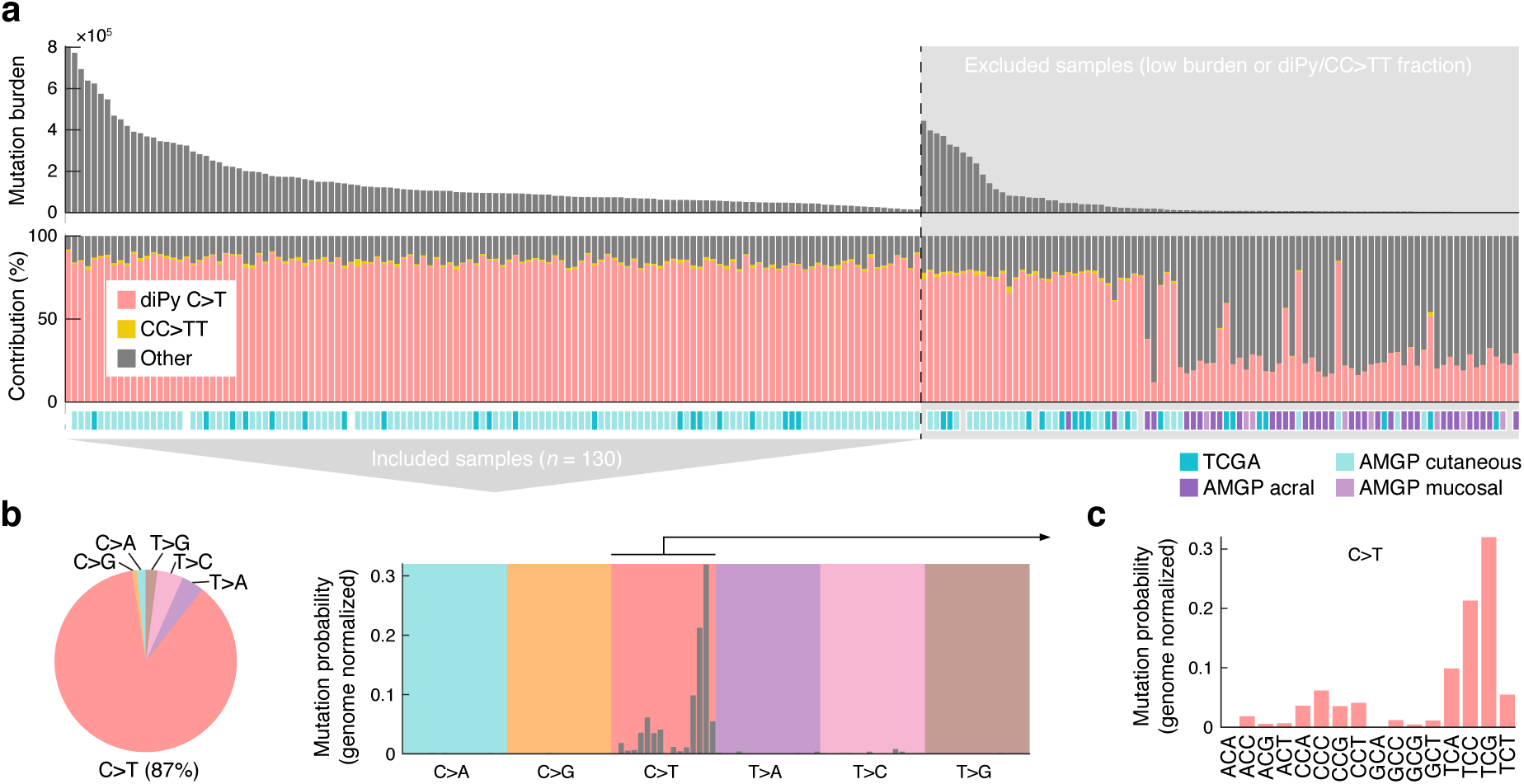
A whole-genome compendium of somatic mutations predominantly induced by UV light. (**a**) Whole genome somatic mutation data from 221 melanomas was initially assembled from earlier studies^16,17^ and a subset of 130 samples with high burden (≥10,000 mutations) and with a high fraction (≥80%) of mutations having characteristics of UV photoproduct formation (C>T in a dipyrimidine context or CC>TT) were included for further study. TCGA, The Cancer Genome Atlas; AMPG, Australian Melanoma Genome Project. (**b**) SNVs (*n* = 19.7×10^6^) in the final dataset are predominantly C>T. (**c**) Trinucleotide signature (genome normalized) for included SNVs show mutations primarily at dipyrimidines, characteristic of mutations arising from UV photoproduct formation.

Expectedly, single nucleotide variants (SNVs) in the resulting dataset were predominantly C>T (87%) with a trinucleotide signature closely resembling the canonical UV signature (“Signature 7”^5^), where T<u>C</u>G and T<u>C</u>C (mutated base underscored) has the highest mutation probabilities after normalization for genomic trinucleotide frequencies (**Fig. 1b-c**). 99.5% of C>T mutations, which we give particular focus in this study, were in dipyrimidine contexts, consistent with the vast majority being canonical UV photoproducts mutations. This established a large (19.7×10^6^ SNVs) compendium of primarily UV-induced mutations to facilitate further study of their sequence properties.

### Promoter-related chromatin states exhibit a unique UV trinucleotide signature

While it is known that the relative contributions of mutational processes may vary across genomic features^18^, less is known about genomic variability in the characteristics of a single process. To this end, we investigated how the trinucleotide signature, normalized by local sequence composition, varied across chromatin states in the present UV-dominated mutational dataset.

Based on a segmentation of the genome into 15 chromatin states (ChromHMM^19^ model based on RoadMap epigenomic data^20^), we found that while all genomic regions exhibited a general UV-like dipyrimidine-related trinucleotide signature, there was also notable variability (**Supplementary Fig. 1**). Specifically, principal components analysis (PCA) appeared to separate the signatures based on whether the corresponding regions were related to transcription start sites (TSSs)/promoters, with E1 (“Active TSS”) and E15 (“Quiescent/low”) representing opposite extremes along the first component (**Fig. 2a**). Further, non-TSS-related states, encompassing the vast majority (98.7%) of the genome, showed strong similarity to the canonical UV “Signature 7”^5^ while TSS-related regions deviated (**Fig. 2b**).

**Figure 2.**
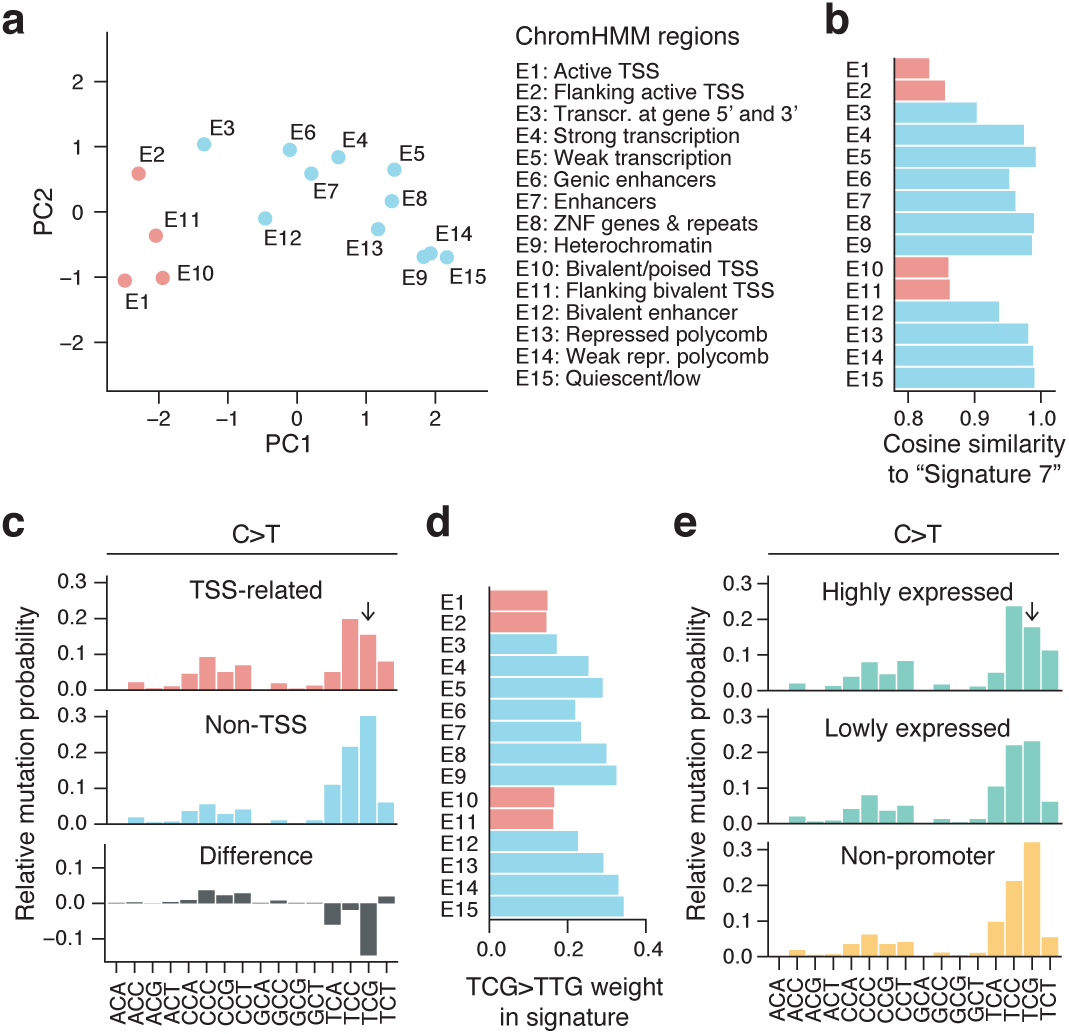
Variation in the UV trinucleotide signature across chromatin states. (**a**) PCA plot of trinucleotide signatures across 15 ChromHMM chromatin states^19^. Transcription start site (TSS)-related regions are indicated in red. (**b**) Similarity (cosine) to the canonical UV signature, “Signature 7”^5^, for each genomic region. (**c**) Pooled C>T trinucleotide signature (local genome normalized) for the TSS-related regions (E1, E2, E10 and E11; red) and remaining non-TSS regions (blue), revealing a reduced mutation rate at TCG in the former. The difference between the two is shown in gray. Frequencies are normalized to sum to 1. (**d**) The C>T substitution frequency at T<u>C</u>G (weight in normalized signature) varies across ChromHMM regions and is reduced in TSS-related states. (**e**) Trinucleotide signature in promoters (500 bp upstream regions) of highly (top 25%) compared to lowly (bottom 25%) expressed genes, with TTC>TCG substitution rate being more reduced in the former category. The two sets were selected from 20,017 annotated coding genes.

Examination of the signature in TSS-related regions revealed that C>T substitutions in the T<u>C</u>G context were notably reduced (**Fig. 2c-d**). TCG, while relatively infrequent in the genome, normally has the highest weight (highest probability of being mutated) in the normalized UV trinucleotide signature (**Fig. 1c**). This may be due to facilitated CPD formation at 5-methylcytosines^21,22^ (5mC), prevalent at CpGs throughout the genome but not in promoters^23^, thus possibly also explaining the deviating signature. Consistent with this model, a recent analysis of *XPC* -/- cutaneous squamous cell carcinomas (cSCC) lacking global nucleotide excision repair (NER), which repairs UV photoproducts, revealed a generally reduced mutation burden in promoters explained by a reduction in Y<u>C</u>G mutations (Y = C/T) that varied with methylation level^24^. C<u>C</u>G mutations, which should theoretically be similarly affected, were slightly increased rather than reduced (**Fig. 2c**), plausibly explained by its low frequency in combination with signature weights being relative rather than absolute.

In agreement with the above results, we found that promoters of annotated genes (500 bp upstream regions) displayed a similar reduction at T<u>C</u>G, which was more pronounced for highly compared to lowly expressed genes (**Fig. 2e**). We conclude that promoter regions show a unique UV trinucleotide signature dominated by TCC>TTC rather than TCG>TTG, possibly explained by reduced CPD formation due to decreased CpG methylation in promoters.

### Local UV trinucleotide signature varies with methylation level

Next, we further investigated the relationship between trinucleotide signature and methylation levels using available whole-genome bisulfite sequencing data^20^. CpG methylation was frequent in non-TSS chromatin states (82.5% combined) but heavily reduced in TSS-related regions (11.2% in E1, “Active TSS”; **Fig. 3a**). Consequently, the weight for TCG>TTG in the signature correlated positively with methylation level across regions (**Fig. 3b**; Pearson’s *r* = 0.82, *P* = 1.8×10^−4^).

**Figure 3.**
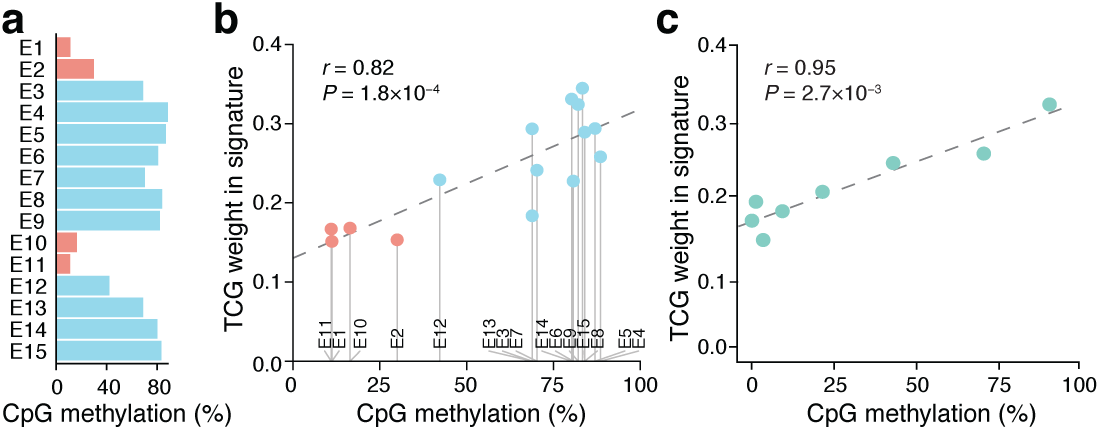
UV trinucleotide signature in promoters vary with methylation. (**a**) Extensive variability in CpG methylation across ChromHMM chromatin states. (**b**) Positive correlation between methylation level and the weight of TCG>TTG substitutions in the trinucleotide signature (local genome normalized) across chromatin states. (**c**) The weight of TCG>TTG substitutions in the trinucleotide signature of promoters varies positively with methylation level. 12,239 annotated coding gene promoters with sufficient bisulfite coverage were divided into 10 methylation level bins (the first three, all representing 0%, were merged into one; x-axis indicates average methylation). Pearson’s correlation coefficients across regions/bins are indicated in panels **b** and **c**.

In annotated promoters, representing a more homogenous set compared to the ChromHMM regions, the weight for TCG>TTG in the signature correlated strongly with increasing methylation level (**Fig. 3c**; Pearson’s *r* = 0.95, *P* = 2.7×10^−3^). These results further solidify a relationship between the deviating UV trinucleotide signature in promoters and reduced methylation in these regions.

### Reduced pyrimidine dimer formation by UVB in promoters with reduced CpG methylation

Given the correlation between methylation level and UV signature characteristics, we next sought to directly determine whether this may be explained by differential DNA damage formation at CpGs. Notably, it has been shown that 5mC facilitates CPD formation by UVB (280-315 nm), the main inducer of CPDs in sunlight, but not by UVC (100-280 nm)^25,26^, which does not penetrate the atmosphere. Despite this, genome-wide studies of CPD formation in human cells to date have all been performed using UVC^14,15,27,28^.

To address this, we mapped CPDs genome-wide in human A375 melanoma cells immediately following exposure to UVB (310 nm), using a protocol based on T4 endonuclease V digestion and Illumina sequencing as described previously for UVC^15^ (**Fig. 4a**). Two independent maps were generated, and previously published CPD data for UVC (254 nm), generated by the same protocol, were included for comparison^15^. CPDs were preferably detected at TT, TC, CT and CC dinucleotides as expected, notably with lower TT and higher CC frequencies relative to UVC, further supporting that the two wavelength ranges are not physiologically equivalent (**Fig. 4b**). Interestingly, an elevation at CG, weak compared to the dipyrimidines but observed in all conditions including non-UV controls, was found to be methylation-dependent, suggesting occasional T4 endonuclease V cleavage at methylated CG dinucleotides (**Supplementary Fig. 2**). In total 77.1 million UVB-induced CPDs were mapped to dipyrimidines throughout the genome and used for further analyses.

**Figure 4.**
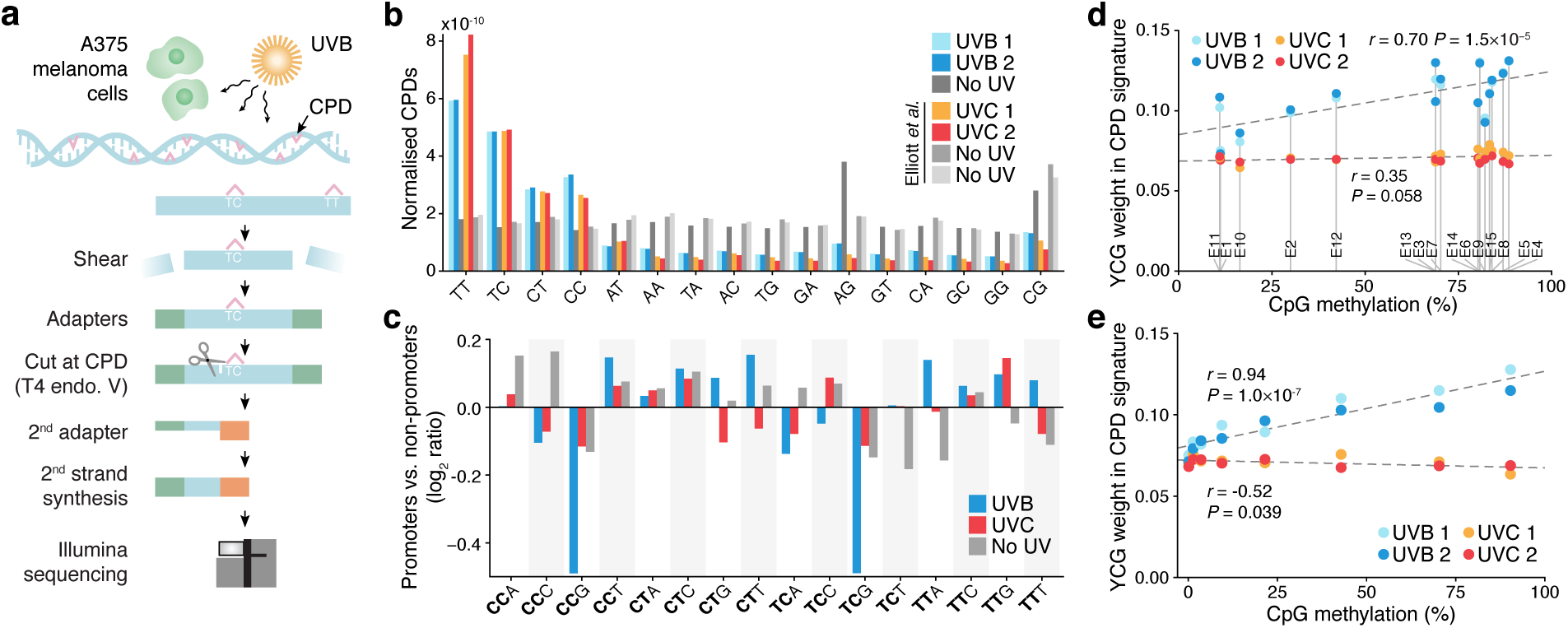
Genome-wide mapping of UVB-induced pyrimidine dimers reveals reduced DNA damage in promoters with reduced CpG methylation. (**a**) Simplified protocol overview for genome-wide mapping of CPDs induced by UVB in A375 human melanoma cells. Cells were treated with 10,000 J/m^2^ UVB (310 nm) and DNA was harvested immediately for analysis. (**b**) Genome-wide CPDs counts for all dinucleotides, normalized with respect to genomic dinucleotide counts and library size, showing preferable detection at dipyrimidines as expected. UVC results from Elliott, et al. ^15^ were included for comparison. (**c**) Reduced UVB-induced DNA damage at <u>YC</u>G sites in promoters. Comparison of the CPD trinucleotide signature (relative formation frequency per genomic site) in highly expressed promoters compared to non-promoter regions, expressed as a log2 ratio. Examined patterns include one additional 3’ base following the CPD-forming dipyrimidine (bold) in all possible combinations, to enable comparison between CpG- and non-CpG-adjacent sites. Results are shown for UVB, UVC and no UV controls. (**d**) CPD frequency at <u>YC</u>G (weight in CPD trinucleotide signature) increases with increasing CpG methylation across ChromHMM regions, specifically for UVB. (**e**) CPD frequency at <u>YC</u>G increases with increasing CpG methylation across annotated promoters, specifically for UVB. Bins were defined as in **Fig. 3c**. Pearson’s correlation coefficients across regions/bins are indicated in panels **d** and **e**.

To compare CpG- and non-CpG-adjacent dipyrimidines in terms of CPD formation, we determined “CPD trinucleotide signatures” describing relative CPD frequencies across patterns consisting of a CPD-forming dipyrimidine plus one additional 3’ base. Genome-wide, these signatures differed markedly between UVB and UVC, as expected given the differences in dinucleotide distribution (**Supplementary Fig. 3**).

Next, we compared highly expressed promoters to non-promoters in terms of CPD signature. Importantly, the former were characterized by reduced CPD formation at <u>YC</u>G (CPD underscored) specifically for UVB but not UVC, thus confirming reduced UVB-induced DNA damage at CpGs in promoters (**Fig. 4c**). A similar UVB-specific pattern was observed when comparing highly to lowly methylated genomic regions (**Supplementary Fig. 2**). Likewise, across ChromHMM regions, the general trend was for UVB CPD frequency at <u>YC</u>G to increase with increasing methylation (*r* = 0.70, *P* = 1.5×10^−5^; **Fig. 4d**). Finally, across annotated promoters there was a strong correlation between methylation level and <u>YC</u>G CPD formation specifically by UVB (*r* = 0.94, *P* = 1.0×10^−7^; **Fig. 4e**). These results show that UVB-induced DNA damage at CpGs is reduced in promoters with reduced methylation, consistent with their deviating mutational signature.

### Improved UV signature modeling by addition of longer contextual patterns

While trinucleotide patterns are informative of whether a UV photoproduct can form, thus providing key information regarding mutation probability, it is also clear that longer motifs may be important^9-15^. These signals may in part be detectable by simply considering a larger number of patterns, or using a position weight matrix still centered at the position of interest^9^, but this will obscure longer patterns occurring at highly asymmetric or variable positions relative to the mutation^12-15^. To address this, we devised an extended mutational signature model that considers the central trinucleotide as well presence/absence of longer pentamer patterns at flexible locations within a +/- 10 bp context around a given position, here focusing on the predominant C>T mutations (**Fig. 5a**).

**Figure 5.**
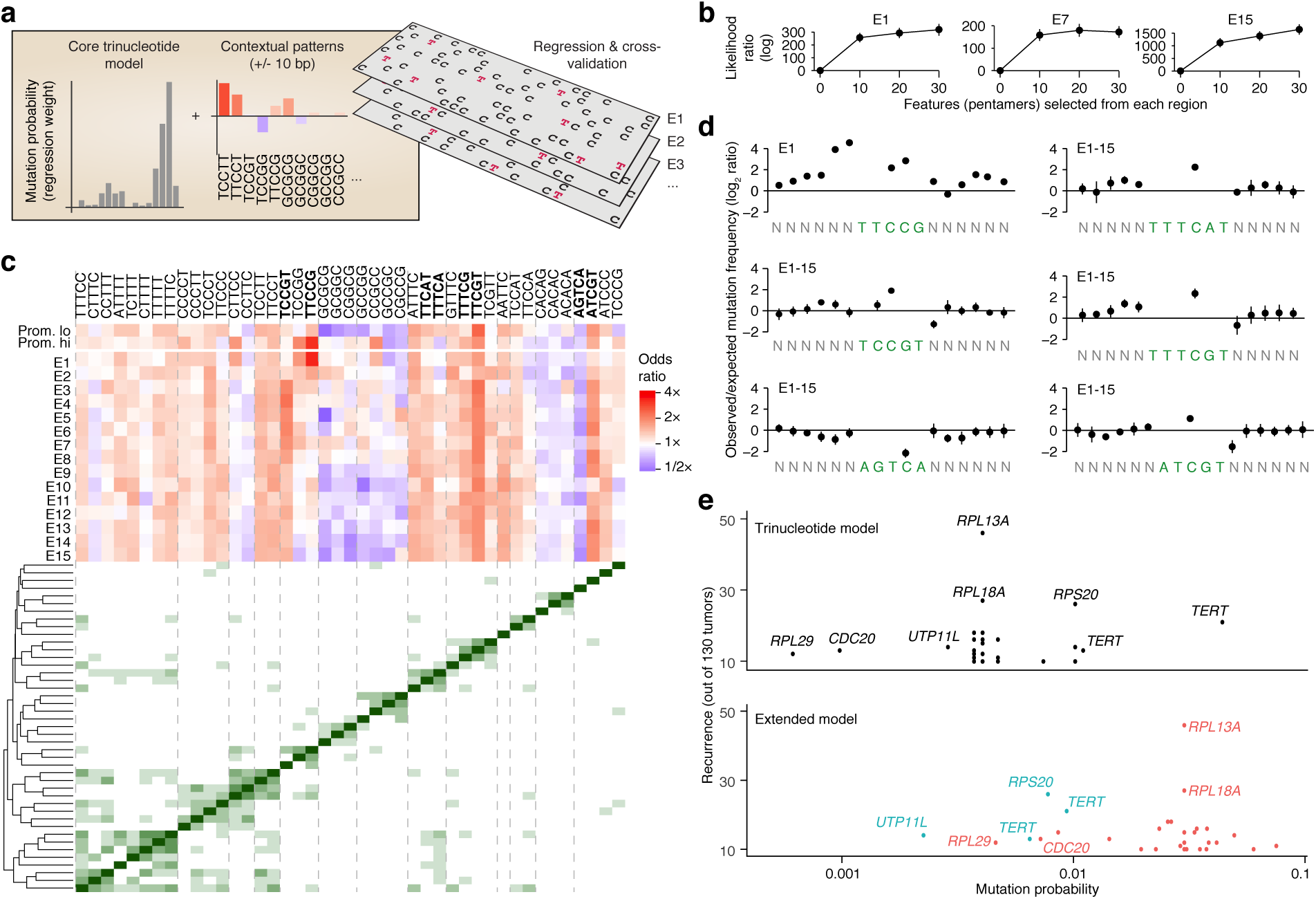
A regression-based signature model reveals extended patterns that are informative of UV mutagenesis in addition to trinucleotides. (**a**) UV mutations (C>T subset) were modeled using logistic regression, taking into account standard trinucleotide patterns as well as presence/absence of longer pentamer patterns occurring anywhere within +/- 10 bp of a given position. Signature models were built repeatedly for each the 15 ChromHMM regions as well promoters (high/low expression), based on 0.5 Mb randomly sampled positions from each region and using a common set of pentamer features (see **Methods**). (**b**) Modeling of observed mutations is improved when long features are considered (all regions shown in **Supplementary Fig. 5**). Ten 0.5 Mb random subsets were evaluated for each region. Log likelihood ratios relative a standard trinucleotide model (zero long features) are shown (bars indicate SD). (**c**) Top heatmap: influence (odds ratio) of different pentamer patterns on mutation probability (blue = stimulatory; red = attenuating) across interrogated regions. Pentamers with low regression weights were excluded for visualization, leaving 43/61 patterns included during feature selection (union of top 20 patterns from each ChromHMM region; see **Methods**). Bottom distance matrix and clustering dendrogram: co-occurrence patterns linking pentamers together into longer motifs. Dashed lines delineate notable clusters. Bold mark patterns highlighted in panel **d**. (**d**) Positional distribution of mutations across select patterns from panel **a** (either individual pentamers or aggregated from multiple pentamers forming a longer consensus motif, as indicated by clustering). Frequencies were normalized to trinucleotide-based expectations given by the underlying sequences. (**e**) Probability of mutagenesis at promoter mutation hotspots (recurrent bases within 500 bp upstream of a TSS) in melanoma, as given by a simple trinucleotide model (upper graph) or the extended model (trinucleotide core model plus longer patterns; lower graph). Locally derived models from corresponding ChromHMM regions were used for all mutations. Recurrence is indicated on the y-axis (*n* ≥ 10). Colors indicate whether probabilities are up (red) or down (blue) in the extended compared to the trinucleotide model.

Extended signatures were determined for each of the 15 ChromHMM regions as well as promoters (highly or lowly expressed) using a common set of candidate pentamer features, briefly by feature selection and fitting of a logistic regression model to randomly subsampled positions from each region (see **Methods**). In principle, this approach allows longer patterns to have stimulating as well as attenuating effects. Cross-validation, again based on random subsampling, showed that the addition of longer motifs consistently led to improved modeling of observed mutations compared to a regular trinucleotide model, which is equivalent to setting the number of long features to zero in the extended model (**Fig. 5b** and **Supplementary Fig. 4**).

The majority of uncovered motifs (pentamers or longer consensus patterns indicated by co-occurrence clustering of pentamers) were associated with increased mutation probability (odds ratio > 1), and the effects were often but not always consistent across the interrogated regions (**Fig. 5c**). TTCCG was detected specifically in promoter-related regions, with E1 (“Active TSS”) and annotated active promoters in particular showing strong positive odds ratios (4.1 and 2.2, respectively, the strongest of all patterns). Elevated mutation frequency was observed in particular at bases preceding TTCCG (**Fig. 5d**). These observations are consistent with a known influence from ETS transcription factor binding site sequences (TTCCG) on CPD formation efficacy, specifically in promoter regions and preferably one or two bases upstream of the motif^12-15^.

In agreement with a demonstrated positive influence from flanking thymines^9-11^, we found patterns related to TTT<u>C</u>NT (main position with elevated mutation rate underscored), as well as other poly-T-containing motifs, to have positive weights across all regions (**Fig. 5c**; exemplified by TTTCAT and TTTCGT in **Fig. 5d**). GC-rich patterns were generally informative of reduced mutation rate, but this effect was notably absent in active promoter regions (**Fig. 5c**). Conceivably, this may be explained by reduced power to call somatic mutations in GC-rich regions^29^, possibly counteracted by a general increase in UV mutation rate at active GC-rich promoters^24,30^. Notable and consistent positive and negative coefficients were also seen for ATCGT and AGTCA, respectively (**Fig. 5c-d**).

Assessment of signals of selection in cancer involves determination of expected mutation rates/probabilities, which may be improved by considering mutational signatures^4^. When applied to recurrent promoter mutations, common in melanoma^31,32^, we found that a regular trinucleotide model failed to give higher mutation probability estimates for non-TERT recurrent sites, generally believed to be passengers^12–15^, compared to *TERT* promoter mutations (C228T and C250T)^33,34^, which are established drivers (**Fig. 5e**). In contrast, the extended model typically gave considerably higher probabilities for the non-TERT sites compared to the *TERT* mutations (**Fig. 5e**), primarily driven by TTCCG (ETS) elements among the former. Exceptions included *UTP11L* and *RPS20*, both lacking ETS motifs, but where C>T mutations in the *RPS20* promoter generate a *de novo* ETS site, similar to *TERT*^33,34^. Low estimated probability for an ETS-related recurrent site in the *RPL29* promoter was explained by counteraction from an uncommon trinucleotide context (A<u>C</u>T). Our results support that consideration of longer contextual patterns in addition to trinucleotides improves modeling of UV mutations, which is beneficial when assessing recurrent mutations in cancer. Moreover, analogous to our observations regarding trinucleotides, this extended UV signature varies between chromatin states.

## DISCUSSION

Earlier studies have revealed a range of mutational signatures representative of different mutational processes, and have shown that the activities of these processes vary between tumors, cancer types and genomic features^1,18^. Here, by detailed characterization of mutations induced by UV radiation through pyrimidine dimer formation, we highlight an additional layer of complexity in the form of intragenomic heterogeneity in the trinucleotide signature characteristics of a single process. Further, we find that longer contextual patterns, which were here combined with trinucleotides into an extended signature model, are informative of mutation probability. Notably, intragenomic variability is again observed for the extended signature. Signatures are thus not static but may vary depending on genomic context, which may be considered in situations such as driver mutation detection where accurate modeling of mutation rates is important.

We specifically find that the UV trinucleotide signature deviates in promoters, primarily due to reduced T<u>C</u>G mutations, which was linked to methylation levels. A methylation-related reduction in mutations at promoter CpG sites has previously been noted in NER-deficient cSCC tumors, proposedly due to reduced CPD formation^24^, but an impact from this effect on the general UV signature in repair-proficient cells has to our knowledge not previously been described or quantified. By generating the first human genome-wide map of UVB-induced CPDs, we here directly demonstrate that DNA damage is reduced at these sites. Importantly, this is not testable using existing UV damage maps generated using UVC^25,26^, and our results support marked physiological differences between the two wavelength ranges. UVC does not penetrate the atmosphere, and previous UV damage data thus fails to accurately reflect sunlight-induced DNA damage patterns in tumors.

Incorporation of longer patterns into a signature model led to improved modelling of observed UV mutations, and our results suggest that this may be beneficial in the context of methods for evaluating signals of selection in cancer. More work is needed for full comprehension of the mechanistic basis of some of the uncovered sequence motifs. Finally, it can be noted that the analyses in this study were considerably simplified by the purity and abundance of UV mutations in skin cancers, eliminating the need for deconvolution strategies. A future prospect, requiring further methodological development, is to more broadly address hypotheses regarding intragenomic signature variability and extended sequence patterns, as these questions are relevant also to other cancers and mutational processes.

## MATERIALS AND METHODS

### Whole genome mutation calls

Mutation calls from the Australian Melanoma Genome Project (AMGP) whole genome sequencing cohort^16^ were obtained from the International Cancer Genome Consortium’s (ICGC) database^35^. These data were pooled with mutation calls from The Cancer Genome Atlas (TCGA) melanoma whole genome cohort^17^, called as described previously^32^. Population variants (dbSNP v138) were removed and in cases where multiple samples from the same patient were available, the sample with the highest median allele frequency was maintained, resulting in a total of 221 tumors. From these, a subset of 130 tumors with heavy UV mutation burden were selected for subsequent analyses (>80% dipyrimidine C>T or CC>TT and total burden >10,000 mutations).

### Gene annotations and ChromHMM genome segmentation data

Gene annotations from GENCODE^36^ v19 were used to define TSS positions for 20,017 uniquely mapped coding genes, disregarding chrM and considering the 5’-most annotated transcripts while excluding non-coding isoforms. Promoters were defined as 500 bp regions upstream of TSSs. Processed RNA-seq data for the TCGA subset of samples were obtained from Ashouri, et al. ^37^, allowing lower and upper expression quartiles to be determined for promoters. ChromHMM^19^ chromatin state genomic segmentations based on epigenomic data from foreskin melanocytes (Roadmap celltype E059, core 15-state model) were obtained from the Roadmap Epigenomics Project (http://www.roadmapepigenomics.org). Positions within 100 bp of a satellite class repeat from RepeatMasker ^38^ were excluded from all regions in all analyses, to avoid erroneously mapped reads in CPD and other datasets.

### CpG methylation analyses

Bisulfite-determined CpG methylation data from leg skin were acquired from ENCODE^39^ (accession ENCFF219GCQ), and coordinates were converted from hg38 to hg19 using liftOver^40^. For methylation analyses of promoters, ChromHMM regions, and genomic bins, methylation levels were defined as the average across all CpGs in that segment, after removing CpGs below a minimum coverage threshold of 5. Segments with fewer than 5 (promoters) or 10 (genomic bins) CpGs were excluded from further analyses, while no such threshold was used for the larger ChromHMM regions. Promoters were initially grouped by methylation level into 10 equally sized bins followed by merging of the lower 3 bins which all represented 0% methylation.

### Genome-wide mapping of UVB-induced cyclobutane pyrimidine dimers

A375 cells were grown in DMEM + 10% FCS + Penicillin/streptomycin (Gibco, Carlsbad, MA) and were treated with 10,000 J/m^2^ UVB 310 nm (7 mins @ 25 J/m^2^/s) using a UV-2 Ultraviolet radiation system (Tyler Research Corporation, Canada) in duplicates (UVB 1 and UVB 2), and DNA from untreated cells was isolated as a control (No UV). Appropriate UVB dose was determined by T4 endonuclease V (NEB) digestion followed by analysis on a 1% alkaline gel (**Supplementary Fig. 5**). CPD sequencing then proceeded as described in Elliott, et al. ^15^. All adapter oligos, including additional indexes used, are indicated in **Supplementary Table 2**. The indexed libraries were pooled and sequenced with a NextSeq 500 using the High Output kit (Illumina, San Diego CA). The data has been deposited in GEO under accession GSE127966. Existing UVC CPD data (UVC1, UVC2, No UV1, No UV2) was obtained from Elliott, et al. ^15^.

### CPD bioinformatics

FastQ files were aligned with Bowtie 2 version 2.3.1^41^ to hg19 with default parameters. Duplicate reads, identified with Picard MarkDuplicates version 2.18.23^42^ with default parameters, were disregarded. For all subsequent analysis, R was used with Bioconductor^43^ packages. CPD positions were defined as two bases upstream on the opposite strand of the first mate in each read pair. CPD trinucleotide signatures were determined by considering patterns consisting of a CPD-forming dinucleotide followed by an additional 3’ base, to enable CpG-related CPDs to be discerned from others. Signature weights were calculated by normalizing CPD counts by local genomic trinucleotide frequencies, followed by scaling of the values such that all shown trinucleotides sums to one. Log2 ratios, used to compare CPD signatures between conditions, were calculated based on these scaled weights.

### Analysis of trinucleotide and extended mutational signatures

Trinucleotide signatures were determined and presented in normalized form throughout the study, dividing observed mutation counts by local genomic trinucleotide counts followed by scaling of signatures weights such that they sum to one. For a given genomic site, the corresponding weight can be interpreted as the relative probability for mutagenesis at this position.

Logistic regression was used to model the impact of longer contextual pentamer patterns, occurring within a +/- 10 bp region of a given position, on mutation probability. These were considered in addition to trinucleotide patterns in the same model. We here focused on the predominant C>T substitutions, but other substitution types can in principle easily be accounted for using multinomial logistic regression. Only cytosine positions were thus considered, taking both strands into account, with a binary response variable indicating whether a mutation was detected or not at each position. The explanatory variables, all binary indicating presence/absence of specific patterns, consisted of all possible trinucleotide contexts plus a limited set of pentamer motifs determined during a feature selection step. It can be noted that, if the number of contextual patterns is set to zero and the resulting regression weights are transformed to frequency space, the resulting signature is equivalent to the genome-normalized trinucleotide signature described above.

To select a common set of long features to be used across all analyzed regions, Fisher’s exact tests was used to test for motifs that were enriched or depleted at mutated positions. 500 kb random subsets of cytosine positions were analyzed this way for each ChromHMM region, and the highest-ranking motifs from each chromHMM region were subsequently pooled. Both pentamers and hexamers were initially evaluated, as well as rank cutoffs of 10, 20 and 30 motifs per region, as indicated in **Fig. 5b**. Based on results from repeated regression and validation on separate random subsets (see below), we opted for pentamers and a rank cutoff of 20 for the final analyses, resulting in a final feature set of 61 pentamers.

Regression models were trained 10 times for each analyzed genomic region, each time on a random 500 kb subset. Each model was evaluated on a separate random subset by determining the log likelihood of the observed data relative to a basic trinucleotide model. The median of the coefficients from the 10 models was used as a consensus model for each region. The same procedure was performed on highly and lowly expressed promoter regions, as defined above. Weak coefficients (log odds within between 0.8 and 1.25 in all region) were excluded during visualization. Remaining motifs were analyzed for co-occurrence across 60 kb of mutated positions (4 kb sampled from each of the 15 ChromHMM regions) using complete linkage hierarchical clustering with Euclidean distance. Select motifs with strong coefficients were further analyzed with respect to positional distribution of mutations. These included pentamers from the model as well consensus hexamers identified manually based on co-occurrence clustering and sequence similarity. For each position, the number of observed mutations were compared against the expected number of mutations based on a trinucleotide model. When comparing the extended signature model to a basic trinucleotide model with respect to estimated mutation probabilities at recurrently mutated positions in promoters (minimum recurrence 10/130 samples), the appropriate extended or trinucleotide model from the corresponding ChromHMM region was used for each site.

## ACKNOWLEDGEMENTS

The results published here are in whole or part based upon data generated by The Cancer Genome Atlas pilot project established by the NCI and NHGRI, as well as ICGC. Information about TCGA and the investigators and institutions who constitute the TCGA research network can be found at “http://cancergenome.nih.gov”. We are most grateful to the patients, investigators, clinicians, technical personnel, and funding bodies who contributed to TCGA and ICGC, thereby making this study possible. We would like to thank the ENCODE consortium and the Richard Myers, HAIB lab for generating and providing the bisulfite methylation data used in this paper. The computations were in part performed on resources provided by SNIC through Uppsala Multidisciplinary Center for Advanced Computational Science (UPPMAX) under project b2012108. This work was supported by grants from the Knut and Alice Wallenberg Foundation (E.L), the Swedish Foundation for Strategic Research (E.L), the Wenner-Gren Foundation (E.L), the Swedish Medical Research Council (E.L), the Swedish Cancer Society (E.L) and the Lars Erik Lundberg Foundation for Research and Education (E.L.).

## REFERENCES

1. Alexandrov, L.B. & Stratton, M.R. Mutational signatures: the patterns of somatic mutations hidden in cancer genomes. Curr Opin Genet Dev 24, 52–60 (2014).

2. Secrier, M. et al. Mutational signatures in esophageal adenocarcinoma define etiologically distinct subgroups with therapeutic relevance. Nat Genet 48, 1131–41 (2016).

3. Ma, J., Setton, J., Lee, N.Y., Riaz, N. & Powell, S.N. The therapeutic significance of mutational signatures from DNA repair deficiency in cancer. Nat Commun 9, 3292 (2018).

4. Ding, L., Wendl, M.C., McMichael, J.F. & Raphael, B.J. Expanding the computational toolbox for mining cancer genomes. Nat Rev Genet 15, 556–70 (2014).

5. Alexandrov, L. et al. Signatures of mutational processes in human cancer. Nature 500, 415–421 (2013).

6. Lawrence, M.S. et al. Mutational heterogeneity in cancer and the search for new cancer-associated genes. Nature 499, 214–8 (2013).

7. Helleday, T., Eshtad, S. & Nik-Zainal, S. Mechanisms underlying mutational signatures in human cancers. Nat Rev Genet 15, 585–98 (2014).

8. Ikehata, H. & Ono, T. The mechanisms of UV mutagenesis. J Radiat Res 52, 115–25 (2011).

9. Krauthammer, M. et al. Exome sequencing identifies recurrent somatic RAC1 mutations in melanoma. Nat Genet 44, 1006–14 (2012).

10. Brash, D.E. & Haseltine, W.A. UV-induced mutation hotspots occur at DNA damage hotspots. Nature 298, 189–92 (1982).

11. Chung, L.H. & Murray, V. An extended sequence specificity for UV-induced DNA damage. J Photochem Photobiol B 178, 133–142 (2018).

12. Fredriksson, N.J. et al. Recurrent promoter mutations in melanoma are defined by an extended context-specific mutational signature. PLoS Genet 13, e1006773 (2017).

13. Colebatch, A.J. et al. Clustered somatic mutations are frequent in transcription factor binding motifs within proximal promoter regions in melanoma and other cutaneous malignancies. Oncotarget 7, 66569–66585 (2016).

14. Mao, P. et al. ETS transcription factors induce a unique UV damage signature that drives recurrent mutagenesis in melanoma. Nat Commun 9, 2626 (2018).

15. Elliott, K. et al. Elevated pyrimidine dimer formation at distinct genomic bases underlies promoter mutation hotspots in UV-exposed cancers. PLoS Genet 14, e1007849 (2018).

16. Hayward, N.K. et al. Whole-genome landscapes of major melanoma subtypes. Nature 545, 175 (2017).

17. The Cancer Genome Atlas Network. Genomic Classification of Cutaneous Melanoma. Cell 161, 1681–1696 (2015).

18. Morganella, S. et al. The topography of mutational processes in breast cancer genomes. Nat Commun 7, 11383 (2016).

19. Ernst, J. & Kellis, M. Chrom HMM: automatingchromatin-statediscoveryand characterization. Nat Methods 9, 215–6 (2012).

20. Roadmap Epigenomics, C. et al. Integrative analysis of 111 reference human epigenomes. Nature 518, 317–30 (2015).

21. Drouin, R. & Therrien, J.P. UVB-induced cyclobutane pyrimidine dimer frequency correlates with skin cancer mutational hotspots in p53. Photochem Photobiol 66, 719–26 (1997).

22. Tommasi, S., Denissenko, M.F. & Pfeifer, G.P. Sunlight induces pyrimidine dimers preferentially at 5-methylcytosine bases. Cancer Res 57, 4727–30 (1997).

23. Jones, P.A. Functions of DNA methylation: islands, start sites, gene bodies and beyond. Nat Rev Genet 13, 484–92 (2012).

24. Perera, D. et al. Differential DNA repair underlies mutation hotspots at active promoters in cancer genomes. Nature 532, 259–63 (2016).

25. Mitchell, D.L. Effects of cytosine methylation on pyrimidine dimer formation in DNA. Photochem Photobiol 71, 162–5 (2000).

26. Rochette, P.J. et al. Influence of cytosine methylation on ultraviolet-induced cyclobutane pyrimidine dimer formation in genomic DNA. Mutat Res 665, 7–13 (2009).

27. Hu, J., Adebali, O., Adar, S. & Sancar, A. Dynamic maps of UV damage formation and repair for the human genome. Proc Natl Acad Sci U S A 114, 6758–6763 (2017).

28. Garcia-Nieto, P.E. et al. Carcinogen susceptibility is regulated by genome architecture and predicts cancer mutagenesis. EMBO J 36, 2829–2843 (2017).

29. Rheinbay, E. et al. Recurrent and functional regulatory mutations in breast cancer. Nature 547, 55–60 (2017).

30. Sabarinathan, R., Mularoni, L., Deu-Pons, J., Gonzalez-Perez, A. & Lopez-Bigas, N. Nucleotide excision repair is impaired by binding of transcription factors to DNA. Nature 532, 264–7 (2016).

31. Melton, C., Reuter, J.A., Spacek, D.V. & Snyder, M. Recurrent somatic mutations in regulatory regions of human cancer genomes. Nat Genet 47, 710–6 (2015).

32. Fredriksson, N.J., Ny, L., Nilsson, J.A. & Larsson, E. Systematic analysis of noncoding somatic mutations and gene expression alterations across 14 tumor types. Nat Genet 46, 1258–63 (2014).

33. Horn, S. et al. TERT promoter mutations in familial and sporadic melanoma. Science 339, 959–61 (2013).

34. Huang, F.W. et al. Highly recurrent TERT promoter mutations in human melanoma. Science 339, 957–9 (2013).

35. The International Cancer Genome Consortium. International network of cancer genome projects. Nature 464, 993 (2010).

36. Derrien, T. et al. The GENCODE v7 catalog of human long noncoding RNAs: analysis of their gene structure, evolution, and expression. Genome Res 22, 1775–89 (2012).

37. Ashouri, A. et al. Pan-cancer transcriptomic analysis associates long non-coding RNAs with key mutational driver events. Nat Commun (2016).

38. Smit, A.F.A., Hubley, R. & Green, P. RepeatMasker Open-4.0. (2013-2015).

39. Dunham, I. et al. An integrated encyclopedia of DNA elements in the human genome. Nature 489, 57–74 (2012).

40. Hinrichs, A.S. et al. The UCSC Genome Browser Database: update 2006. Nucleic Acids Res 34, D590–8 (2006).

41. Langmead, B. & Salzberg, S.L. Fast gapped-read alignment with Bowtie 2. Nature methods 9, 357–359 (2012).

42. Broad Institute. Picard Tools. Version 2.18.7 edn (2018).

43. Huber, W. et al. Orchestrating high-throughput genomic analysis with Bioconductor. Nat Methods 12, 115–21 (2015).

