## Supplementary figures and tables for "Intragenomic variability and extended sequence patterns in the mutational signature of ultraviolet light"

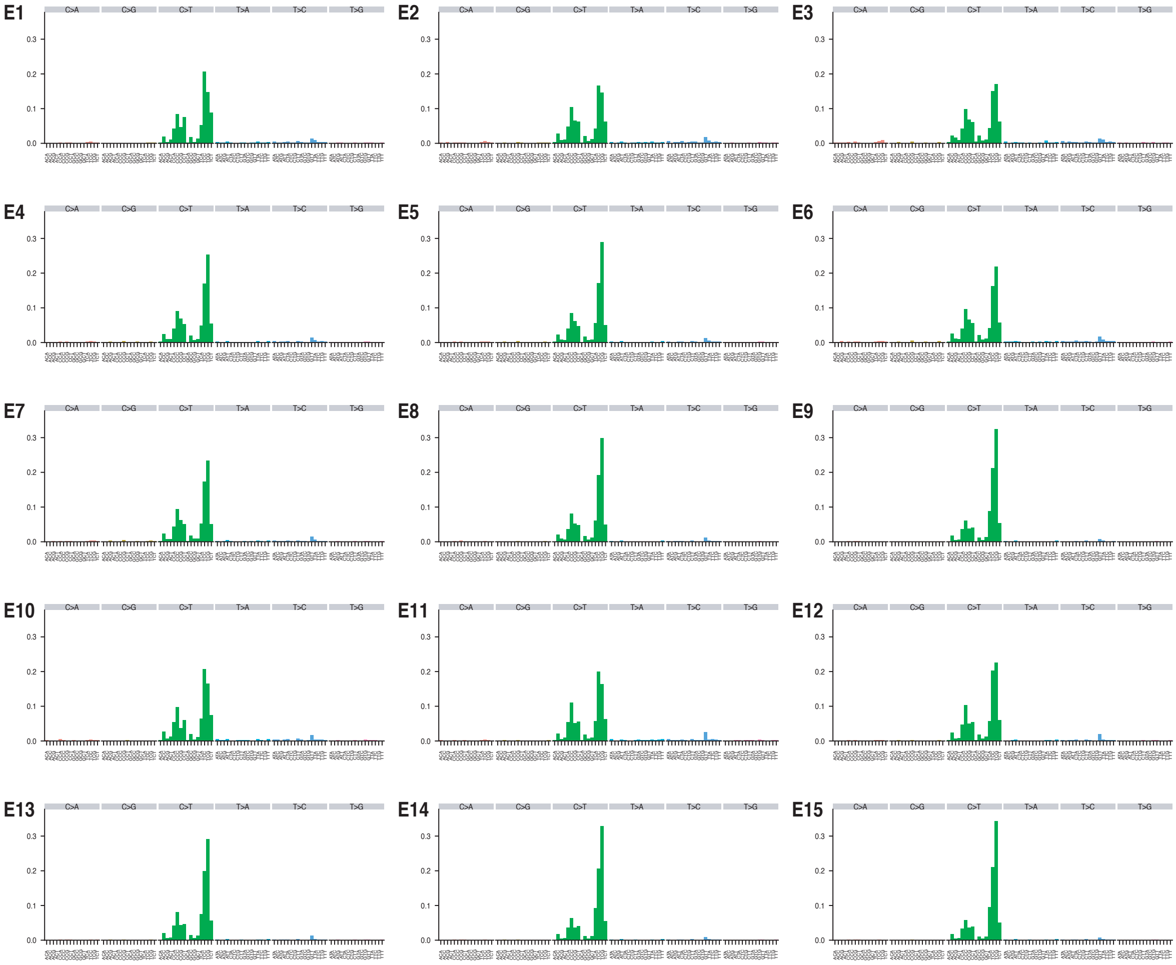


**Figure S1. UV trinucleotide signature in different ChrommHMM chromatin states.** The signatures were normalized for variable trinucleotide sequence content in the respective regions and further normalized to sum up to one. Each bar thus shows the probability of mutagenesis at a given trinucleotide in relative terms, compared to other trinucleotides.

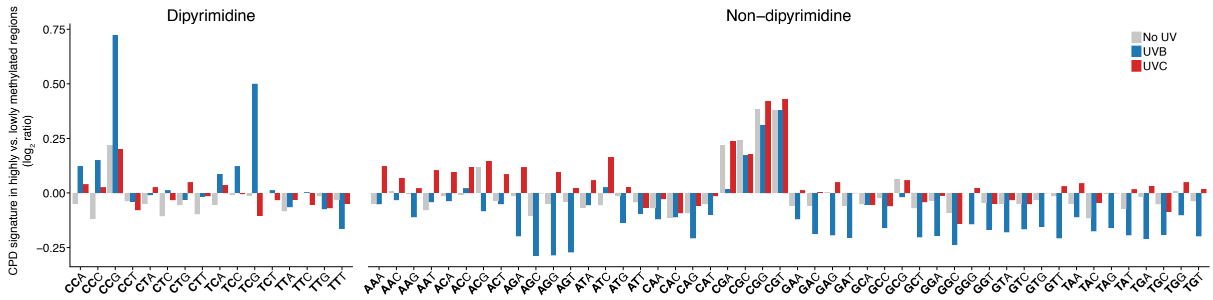


**Figure S2. CPD formation signature in highly vs. lowly methylated genomic regions.** The CPD trinucleotide signature (relative formation frequency per genomic site) in highly (>80% CpG methylation) vs. lowly (>20% CpG methylation) methylated regions (1 kb genomic bins) were compared by means of a $\log_2$ ratio, confirming methylation-dependent elevation in CPD formation at CpG-flanking dipyrimidines specifically in response to UVB (blue). Examined patterns include all trinucleotides, where the first two bases (bold) represent the position at which a CPD was detected. Notably, CPD detections at CG dinucleotides, which cannot form CPDs and thus represent false positives, were found to be methylation-dependent. Although frequencies for these events were low compared to actual CPD-forming dipyrimidines (**Fig. 4**), such detections were observed in all conditions including no-UV-controls, suggesting occasional cleavage by T4 endonuclease V at methylated CpGs independently of CPD formation. Signatures were normalized with respect to genomic sequence content in the respective regions and further normalized to sum to one. Trinucleotides are presented in alphabetical order, separated by CPD-forming and non-CPD-forming patterns. Results for UVB, UVC and no UV controls, all pooled, are shown as separate bars.

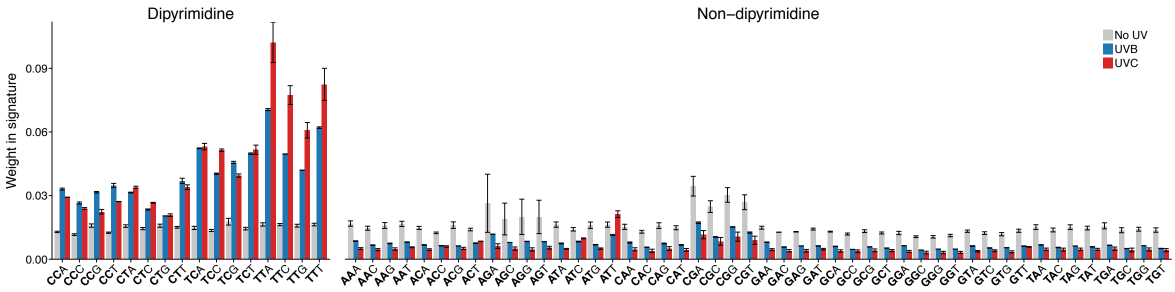


**Figure S3. Genome-wide CPD trinucleotide signature for UVB and UVC.** Examined patterns include all trinucleotides, where the first two bases (bold) represent the position at which a CPD was detected. Signatures were normalized with respect to genomic sequence content in the respective regions and further normalized to sum to one. Trinucleotides are presented in alphabetical order, separated by CPD-forming and non-CPD-forming patterns. Results for UVB, UVC and no UV controls, are shown, and error bars indicate SD ($n$ = 2).

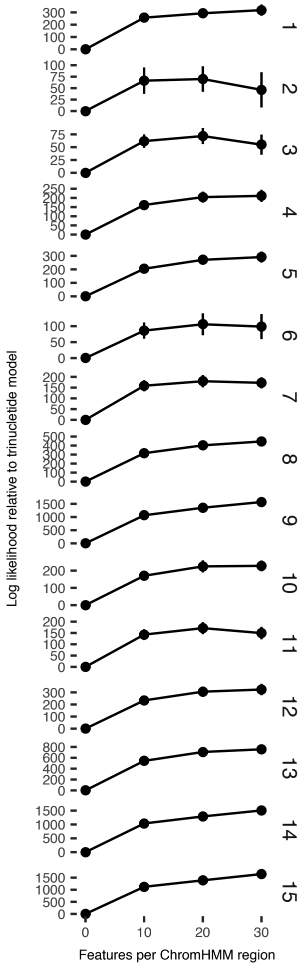


**Figure S4. Likelihood of observed mutation data for the extended signature model as a function of the number of long pentamer features.** The x-axis indicates the number of pentamer features contributed from each ChromHMM region during feature selection, which was performed using Fisher's exact test on 500 kb random subsets of cytosine positions in each region. Selected features from each region (top ranking positive or negatively associated motifs) were pooled into one final set used for training models using logistic regression. This was repeated 10 times for each region on separate 500 kb random subsets. For each model, the likelihood was evaluated based on observed mutation data in a separate random 500 kb subset.

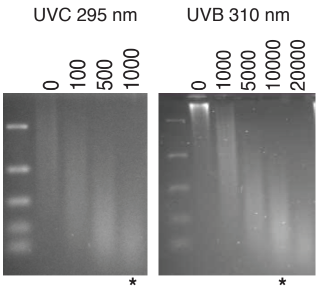


**Figure S5**. **T4 endonuclease digestion of DNA treated with different doses of UV light**. To ensure that CPDs were generated at similar frequencies compared to previously generated UVC data[1], A375 cells were treated with a range of UVC (295nm) and UVB (310 nm) doses. DNA was extracted using the QIAgen Blood Mini kit, and 1 µg of DNA was digested with T4 endonuclease V (NEB). DNA was next purified by phenol/chloroform extraction and ethanol precipitation. DNA was resuspended in alkaline sample buffer and run overnight on a 1% alkaline gel. A UVB dose of 10,000 J/m$^2$ was judged to be appropriate, as indicated by the asterisks.

### SUPPLEMENTARY TABLES

| Primers | Sequence |
|---|---|
| ARC141/142 | 5′-GTGACTGGAGTTCAGACGTGTGCTCTTCCGATCT*T-3′ |
| | 5′-/5Phos/AGATCGGAAGAGCACACGTCTGAACTCCAGTCAC/3AmMO/-3′ |
| ARC143/144 | 5′-/5Biosg/ACACTCTTTCCCTACACGACGCTCTTCCGATCTNNNNNN/3AmMO/-3′ |
| | 5′-/5Phos/AGATCGGAAGAGCGTCGTGTAGGGAAAGAGTGT/3AmMO/-3′ |
| ARC154 | 5′-ACACTCTTTCCCTACACGACGCTCTTCCGATCT-3′ |
| ARC49 | 5′-AATGATACGGCGACCACCGAGATCTACACTCTTTCCCTACACGACGCTCTTCCGATCT-3′ |
| ARC78 | 5′-*CAAGCAGAAGACGGCATACGAGAT***CGTGAT**GTGACTGGAGTTCAGACGTGTGCTCTTCCGATCT-3′ |
| ARC87 | 5′-*CAAGCAGAAGACGGCATACGAGAT***CACTGT**GTGACTGGAGTTCAGACGTGTGCTCTTCCGATCT-3′ |
| ARC88 | 5′-*CAAGCAGAAGACGGCATACGAGAT***ATTGGC**GTGACTGGAGTTCAGACGTGTGCTCTTCCGATCT-3′ |

**Table S2. Oligonucleotide sequences for CPD-seq.** Illumina P5 and *P7* adapters are indicated underlined and italicized respectively, and **indexes** are shown in bold and underline. Oligo 5′ modifications are also indicated. All oligos were from Integrated DNA technologies (Coralville, IA).

### SUPPLEMENTARY REFERENCES

1. Elliott, K. *et al.* Elevated pyrimidine dimer formation at distinct genomic bases underlies promoter mutation hotspots in UV-exposed cancers. *PLoS Genet* **14**, e1007849 (2018).
